# ROLE of inter-related population-level host traits in determining pathogen richness and zoonotic risk

**DOI:** 10.1101/123067

**Authors:** Tim C.D. Lucas, Hilde M. Wilkinson-Herbots, Kate E. Jones

## Abstract

Zoonotic diseases are an increasingly important source of human infectious diseases, and host pathogen richness of reservoir host species is a critical driver of spill-over risk. Population-level traits of hosts such as population size, host density and geographic range size have all been shown to be important determinants of host pathogen richness. However, empirically identifying the independent influences of these traits has proven difficult as many of these traits directly depend on each other. Here we develop a mechanistic, metapopulation, susceptible-infected-recovered model to identify the independent influences of these population-level traits on the ability of a newly evolved pathogen to invade and persist in host populations in the presence of an endemic pathogen. We use bats as a case study as they are highly social and an important source of zoonotic disease. We show that larger populations and group sizes had a greater influence on the chances of pathogen invasion and persistence than increased host density or the number of groups. As anthropogenic change affects these traits to different extents, this increased understanding of how traits independently determine pathogen richness will aid in predicting future zoonotic spill-over risk.

## 1. Introduction

Zoonotic diseases are a major source of human infectious disease [1,2]. Epidemics of emerging, zoonotic diseases pose a major threat to human health and economic development [3,4]. The probability of zoonotic spill-over depends on, amongst other factors, the number of pathogen species carried by reservoir host species (pathogen richness) [5]. Empirical, comparative studies across reservoir host species, suggest that host morphological and life-history traits, such as large body size and longevity, correlate strongly with high pathogen richness [6,7]. However, traits related to reservoir host population biology are also expected to affect disease dynamics and therefore influence pathogen richness. Population-level traits such as increased host density [6,8,9], large geographic range size [6,9,10] and greater population structure (nonrandom interactions between individuals) [10,11] have been shown to correlate with high pathogen richness, although the evidence for a relationship with group size (number of individuals in a social group) has been equivocal in many studies [9,10,12–14]. Population size (total number of individuals), an important population-level trait, has rarely been included in comparative studies, despite its importance in describing epidemiological populations [15].

Collinearity between explanatory variables is a common problem in correlative studies, and this issue is exacerbated when there are clear, causal relationships between explanatory variables. There are two particularly clear relationships between the population-level traits associated with pathogen richness. Firstly, host density, *d*, host population size, *N*, and geographic range size, *a*, are, by definition, linked by *d* = *N/a* (see electronic supplementary material, table S1 for all parameters used) and this relationship has broad empirical support [16]. Secondly, host population size can be decomposed into two components, the number of groups, *m*, and the average size of a group, *n*, with *N* = *mn*. Correlative, comparative studies would be especially poor at identifying which, if any, of these traits causally affect pathogen richness. This lack of discriminatory power is particularly important with respect to global change and its effects on zoonotic disease emergence. Population-level traits such as population size and geographic range size, although interrelated, will respond differently to global change and the response will be species specific. Some host species may suffer geographic range contractions while their density remains constant [17]. Other host species might retain their geographic range but have a depressed population density [18]. Only by knowing which of these interrelated traits control pathogen richness will we be able to predict future changes in pathogen richness.

Mechanistic models provide one method for comparing the importance of intrinsically related traits and can provide a deeper understanding of the system than correlative approaches. Theoretical studies have established that a number of host population-level traits are important for epidemiological dynamics and the maintenance of pathogen richness. In particular, host density, population structure and group size are well established as having central roles in pathogen dynamics [19–21]. A number of studies have found that increased host population structure can promote pathogen coexistence [22–24]. While these studies have examined whether these population-level traits can promote pathogen richness, none have attempted to distinguish which might be the most important. Mechanistic models that try to disentangle the interplay between population-level traits including host density, population size, geographic range size, group size and the number of groups are critically needed.

Here, we use multipathogen models to individually vary host population-level traits. We examine a simple deterministic model to establish whether a newly evolved pathogen can invade into an unstructured population in the presence of strong competition from an endemic pathogen strain. We then examine a stochastic, metapopulation model that was parameterised to broadly mimic wild bat populations. We used bats as a case study as group (colony) size is very variable between bat species and bat colonies are often very stable [25–28]. Furthermore, bats are particularly relevant in the context of zoonotic disease as they are thought to be reservoirs for a number of recent, important outbreaks [29,30]. We examined how the interrelated population-level traits affect the ability of a newly evolved pathogen to invade and persist in a population. We used these simulations to examine two specific hypotheses. First, we investigated whether host population size or density more strongly promotes the invasion of a new pathogen. Secondly, we investigated whether the invasion of a new pathogen is more strongly promoted by group size or the number of groups. We found that population size has a much stronger effect on the invasion of a new pathogen than host density. We also found that increasing population size by increasing group size promotes pathogen invasion much more than increasing population size by increasing the number of groups.

## 2. Methods

### (a). Two pathogen SIR model

We developed a multipathogen, susceptible-infected-recovered (SIR) compartment model to examine the probability that a newly evolved pathogen would invade and persist into a population in the presence of an identical, endemic pathogen. Susceptible individuals were counted in class *S* (figure 1) while infected individuals were counted in classes *I*_1_, *I*_2_ and *I*_12_, being individuals infected with Pathogen 1, Pathogen 2 or both, respectively. Recovered individuals, *R*, were immune to both pathogens, even if they had only been infected by one (i.e. there was complete cross-immunity). Furthermore, recovery from one pathogen moved an individual into the recovered class, even if the individual was coinfected (figure 1). Though our study was restricted to two pathogens, this modelling choice allows the model to be easily expanded to include many pathogens. Our assumption of immediate recovery from all other pathogens is likely to be reasonable [31] as any up-regulation of innate immune response or acquired immunity would affect both pathogens equally. The coinfection rate (the rate at which an infected individual is infected with a second pathogen) was adjusted compared to the infection rate by a factor α. Birth rate (Λ) was set equal to the death rate (*μ*), meaning the population did not systematically increase or decrease. New born individuals entered the susceptible class. Infection and coinfection were assumed not to cause increased mortality as bats show no clinical signs of infection for a number of viruses [32,33].

**Figure 1.**
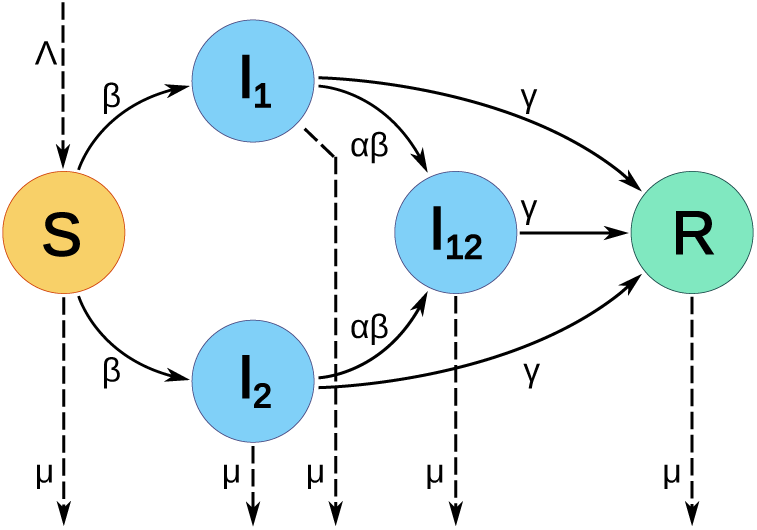
Schematic of the two-pathogen SIR model used. Individuals are in one of five epidemiological classes, susceptible (orange, *S*), infected with Pathogen 1, Pathogen 2 or both (blue, *I*_1_, *I*_2_, *I*_12_, respectively) or recovered and immune from further infection (green, *R*). Transitions between classes occur as indicated by solid arrows and depend on transmission rate (*β*), coinfection adjustment factor (*α*) and recovery rate (*γ*). Births (Λ) and deaths (*μ*) are indicated by dashed arrows. Note that individuals in *I*_12_ move into *R*, not back to *I*_1_ or *I*_2_. That is, recovery from one pathogen causes immediate recovery from the other pathogen.

Population structure is present in bat populations as demonstrated by the existence of subspecies, measurements of genetic dissimilarity and behavioural studies [25,26,34]. Therefore assuming a fully-mixed population is an oversimplification. Consequently, the population was modelled as a metapopulation network with groups being nodes and dispersal between groups being indicated by edges (electronic supplementary material, figure S1). Individuals within a group interacted randomly so that the group was fully mixed. Dispersal occurred at a rate *ξ* between groups connected by an edge in the network. The dispersal rate from a group *y* with degree *k_y_* to group *x* was *ξ*/*k_y_*. Note that this rate was not affected by the degree and size of group *x* and the total rate of dispersal was not affected by the degree of a group. We examined the full model using stochastic, continuous-time simulations, in *R* [35,36]. The full details of the model are given in electronic supplementary material S1.

### (b). Deterministic model

We examined a single-group, deterministic model to establish a baseline expectation for whether a newly evolved pathogen strain could invade into a population in the presence of an identical strain given the assumptions of our two-pathogen SIR model (for details see electronic supplementary material S2). If we first consider the endemic pathogen (Pathogen 1) we have a typical SIR model with vital dynamics [37] with equilibrium values 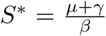 and 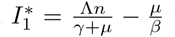 where *β* and *γ* are the transmission and recovery rates and *n* is the group size. When Pathogen 2 is introduced, its rate of change can be written as

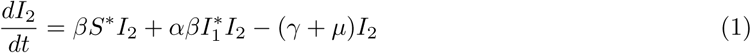
which is greater than zero when *α* (Λ*R*_0_ – μ) *I_2_* > 0 (with *R_0_ =* 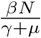 being the basic reproduction number and being equal for the two identical pathogens). As Λ = *μ* due to the assumption of a stable population size, *ΛR*_0_ – *μ* is greater than zero, Therefore, 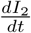 is greater than zero provided α is greater than zero. That is, provided cross-immunity is not complete, Pathogen 2 will invade in this deterministic model. This means that it is only stochastic extinction that would prevent a pathogen from invading into a population. Our more detailed simulations are therefore examining how effectively different population-level traits alleviate stochastic extinction or allow longer term persistence.

### (c). Parameter selection

While some fixed parameters were chosen to approximate those found in wild bat populations, others were chosen for modelling reasons. Assuming equal birth and death rates, Λ and *μ*, were both set to 0.05 per year giving a generation time of 20 years [27,28]. Setting birth and death rates equal gives a stochastically varying population size but given the length of the simulations groups were very unlikely to go extinct. Although it is difficult to directly estimate infection durations in wild bat populations [38], evidence suggests these can be on the scale of days [39] up to months or years [40–42]. Here we set the infection duration, 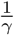, to one year which is a long lasting, but non-chronic, infection. As estimating transmission rates is particularly difficult we used three values of the transmission rate, *β*: 0.1, 0.2 and 0.3. These values were chosen as very high values of *R*_0_ were required so that a reasonable number of simulations experienced an invasion of Pathogen 2. The coinfection adjustment parameter, *α*, was set to 0.1. The deterministic model showed that *α* = 0 and *α* > 0 are qualitatively different with the number of individuals infected with Pathogen 2 being stable and increasing respectively. The case where Pathogen 2 does not invade and spread (*α* = 0) is not important for pathogen richness so we chose a small, non-zero value for *α*. The dispersal rate, *ξ*, was set to 0.01 which yields 17% of individuals dispersing in their lifetime. This relates to a species with juvenile dispersal of a proportion of males and very few females [28,43].

The effect of geographic range size on disease dynamics occurred through changes in the metapopulation network. Dispersal was only allowed to occur between two groups if they were connected nodes in the metapopulation network. The metapopulation network was created for each simulation by randomly placing groups in a square space which represented the geographic range of the species (electronic supplementary material, figure S1). Groups within a threshold distance of each other were connected in the metapopulation network. This meant that the metapopulation network was not necessarily connected (made up of a single connected component). To ensure connected metapopulation networks would have required repeatedly resampling the group locations until a connected metapopulation occurred. However, this would have biased the mean degree, 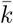. Therefore, it was considered preferential to keep the unconnected networks.

### (d). Experimental setup

A total of 4500 simulations were run. In each simulation, each group in the host population was seeded with 20 individuals infected with Pathogen 1. A ‘burn-in’ of 6 × 10^5^ events was run to allow Pathogen 1 to spread and reach equilibrium. Plotting of preliminary simulations was used to determine that 6 × 10^5^ events was enough to ensure equilibrium. After this burn-in, five host individuals infected with Pathogen 2 were added to one randomly selected group. The invasion and persistence of Pathogen 2 was considered successful if any individuals infected with Pathogen 2 remained at the end of the simulation. As simulations with many individuals and infection events had more events per unit time, the end of the simulation had to be defined in terms of time rather than the number of events. Simulations were run until 75 years had elapsed since the introduction of Pathogen 2. This value was chosen so that pathogens had to persist for multiple host generations in order to be considered persistent.

Three sets of 1500 simulations were run in which three population parameters were varied: (*i*) geographic range size (with corresponding change in host density), (*ii*) group size (with corresponding change in population size), and (*iii*) the number of groups (with corresponding change in population size). To allow comparisons between simulation sets, the parameter that was being varied in each set was assigned its default value multiplied by 0.25, 0.5, 1, 2 and 4. To examine whether host density had a stronger effect on pathogen invasion than population size, results from simulation set *i* were compared to those from sets *ii* and *iii*. To examine whether group size or the number of groups was the more important component of population size, results from simulation set *ii* were compared to those from *iii*.

The spatial scale is arbitrary; only the ratio between the geographic range size and the threshold distance for groups being connected in the metapopulation network had any effect on simulation outcomes. We parameterised the spatial scale by arbitrarily selecting a threshold distance of 100 km˙. The default (10000 km^2^) and upper and lower bounds of the geographic range size (2500–40000 km^2^) were then selected to maximise the range of 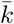 (electronic supplementary material, figure S2) while not having many simulations with networks that were unconnected. That is, we aimed for low host density populations to have relatively sparse population networks, while high host density populations had fully-connected metapopulation networks. This reflects the existence of both isolation-by-distance [43–45] and panmixia [29,46,47] in bat species. The default group size was 400 with a range of 100–1600 which is representative of many bat species [27]. The default number of groups was 20 with a range of 5–80. This minimum value is close to the minimum possible for the population to still be considered a metapopulation, while the maximum value was constrained by computational costs.

In the first set of simulations (*i*), host density was varied by keeping population size constant (*N* = 8000, *n* = 400, *m* = 20) while varying geographic range size. The values of geographic range size used were 2500, 5000, 10000, 20000 and 40000 km^2^ which gave density values of 3.2, 1.6, 0.8, 0.4 and 0.2 animals per km^2^. In the second set of simulations (*ii*), population size was varied by changing group size while the number of groups was kept constant. To keep host density constant, geographic range size was increased as population size increased. The values of group size used were 100, 200, 400, 800 and 1600 while geographic range size was set to 2500, 5000, 10000, 20000 and 40000 km^2^. This gave population sizes of 2000, 4000, 8000, 16000 and 32000 while host density remained at 0.8 hosts per km^2^. In the third set of simulations (*iii*), population size was varied by changing the number of groups while group size was kept constant. Again, to keep host density constant, geographic range size was increased as population size increased. The numbers of groups used were 5, 10, 20, 40 and 80 while geographic range size was set to 2500, 5000, 10000, 20000 and 40000 km^2^. Again, this gave population sizes of 2000, 4000, 8000, 16000 and 32000 while host density remained at 0.8 hosts per km^2^.

### (e). Statistical analysis

For each set of simulations, we fitted binomial GLMs in R [35] with the proportion of invasions of Pathogen 2 as the response variable. To enable comparison between GLMs we divided the explanatory variables by their default values and log_2_ transformed. The explanatory variables for all three sets of simulations therefore became evenly spaced between -2 and 2. To investigate whether an increase in host population size created a stronger increase in probability of pathogen invasion than an equal increase in host density we compared the size (and 95% confidence intervals) of the regression coefficients of host density (*i*) to group size (*ii*) and number of groups (*iii*). To examine whether an increase in group size creates a stronger increase in invasion probability than a proportionally equal increase in number of groups we compared regression coefficients of group size (*ii*) to number of groups (*iii*).

## 3. Results

### (a). Population size and host density

The estimated GLM coefficients were always larger for simulations where population size was varied (sets *ii* and *iii*) than when host density (set *i*) was varied (table 1, electronic supplementary material figure S3). Increasing population size, either by increasing group size or number of groups, increased the probability of invasion (figure 2). The positive relationship between group size and invasion was strong and significant at all transmission rates, while the relationship between number of groups and invasion was weaker but still significant. In contrast, varying host density did not significantly alter invasion probability except for when *β* = 0.3 where there was a significant decrease in invasion probability with increased host density (GLM: coefficient = -0.33, *p* = 0.04). The 95% confidence intervals for the coefficients of group size did not overlap with those for the coefficients of host density at any value of *β* while the 95% confidence intervals for coefficients of number of groups only overlapped with those for host density at *β* = 0.2.

**Table 1.** Regression results comparing effects of group size, number of groups and host density. Estimated regression coefficients and their 95% confidence intervals are from binomial GLM regressions with the proportion of invasions as the response variable and all explanatory variables being standardised by dividing by the default parameter value and applying a log_2_ transform. Group size and number of groups were varied while keeping density equal while density was varied by changing geographic range size while keeping population size equal. Results are given for three transmission (*β*) values.

| $\beta$ | Variable | Coefficient | (95% CI) | p |
| --- | --- | --- | --- | --- |
| 0.1 | Group size | 1.24 | (0.94, 1.59) | $< 10^{-4}$ |
| | Number of groups | 1.03 | (0.43, 1.87) | $3.71 \times 10^{-3}$ |
|  | Density | 0.00 | (-0.41, 0.41) | 1.00 |
| 0.2 | Group size | 2.06 | (1.71, 2.47) | $< 10^{-4}$ |
| | Number of groups | 0.73 | (0.32, 1.24) | $1.54 \times 10^{-3}$ |
|  | Density | 0.13 | (-0.28, 0.55) | 0.54 |
| 0.3 | Group size | 2.69 | (2.23, 3.22) | $< 10^{-4}$ |
| | Number of groups | 0.68 | (0.42, 0.98) | $< 10^{-4}$ |
|  | Density | -0.33 | (-0.66, -0.03) | 0.04 |

**Figure 2.**
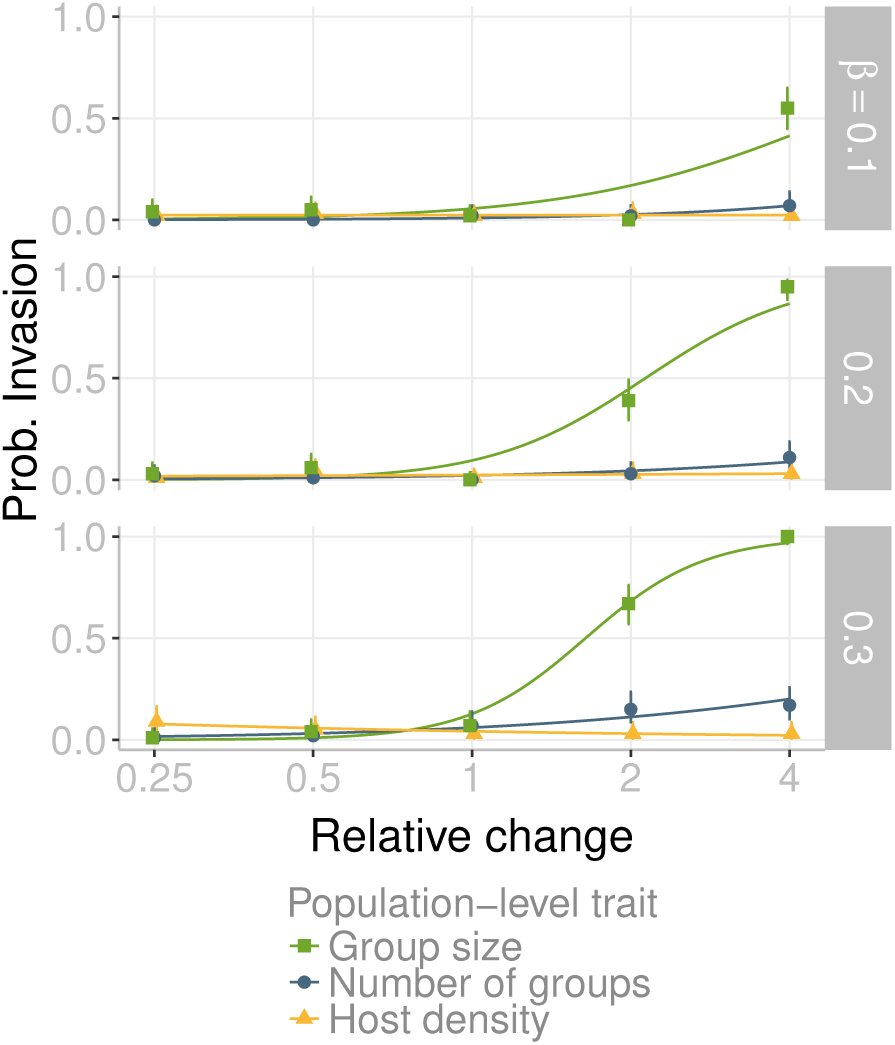
Comparison of the effect of population-level traits on probability of invasion. Population-level traits are group size (green lines, squares), number of groups (blue lines, circles) and host density (yellow lines, triangles). The *x*-axis shows the change (×0.25, 0.5, 1, 2 and 4) in each of these traits relative to the default value. Default values are: number of groups = 20, group size = 400 and host density = 0.8 animals per km^2^. Each point is the mean of 100 simulations and bars are 95% confidence intervals. Each curve was obtained by fitting a binomial GLM. Relationships are shown separately for each transmission value, *β*.

### (b). Group size and number of groups

In all cases, the estimated GLM coefficients were larger for simulations where group size was varied (set *ii*) than when the number of groups (set *iii*) was varied (table 1, electronic supplementary material figures S3 and S4). Increasing either group size or the number of groups increased the probability of invasion but this effect was stronger for group size (figure 2). The 95% confidence intervals of regression coefficients for group size did not overlap with those for number of groups for the simulations with *β* = 0.2 or 0.3 but the confidence intervals did overlap for the simulations when *β* = 0.1.

## 4. Discussion

Overall, our results suggest that population size strongly promotes pathogen richness. In contrast, host density was found to weakly promote pathogen richness if at all. This is in agreement with theoretical arguments that population size is the more natural description of populations [15] but is in contrast to the many comparative studies that find host density to be a significant predictor of pathogen richness [6,8,9]. This suggests that in these comparative studies, host density is acting as a proxy for population size rather than being a causal factor. Alternatively, the local measurements of host density could be indicative of some other trait such as maximum host density, rather than global density. Our results also suggest that a population made up of large groups would have higher pathogen richness than a population made up of many smaller groups. Comparative studies examining group size have had conflicting results [9,10,12,13] with a meta-analysis finding a non-significant relationship between group size and pathogen richness [14]. The largest study specifically on bats found a negative relationship between group size and pathogen richness [13]. Therefore while our results regarding group size are potentially in conflict with the empirical literature, more definitive empirical studies are needed. This conflict may also indicate a difference between pathogen richness accumulation (the focus of this study) and total pathogen richness.

Our results also suggest that host geographic range size does not promote pathogen richness, yet a number of studies have found evidence of this relationship [6,9]. In studies that do not also include population size or host density, geographic range size may be acting as a proxy for population size.

Alternatively this empirical association may be because in wild species, increased host geographic size tends to entail a greater variety of environmental conditions and a greater number of sympatric species which is known to also affect pathogen richness [48] and these factors are not considered in this study. Finally, we found an unexpected negative association between density and invasion probability at *β* = 0.3. Due to the small size of the estimated coefficient, the marginal p-value and the lack of a consistent relationship at other *β* values, we suspect that this is not a real effect but merely due to chance. However, given that in our study, increased density affects disease dynamics by decreasing population structure, this negative relationship does fit with studies that suggest that population structure should allow pathogen coexistence [10,11,23].

Many comparative studies measure population-level traits, yet it is rarely considered how these might be causally related (though statistical correlations between explanatory variables are often handled appropriately). For example, host density is often used in studies [8,9,49], yet host density is directly associated with population size. Our results suggest that despite the association between these two traits, they are not equivalent. These causal relationships between population-level traits should be considered more carefully in future comparative studies. Researchers could include of an interaction term between geographic range size and host density to test for the importance of population size, for example.

This study was limited to one mechanism by which pathogen richness can be increased; the invasion and persistence of a newly evolved pathogen [50]. However, other processes such as pathogen extinction are also likely to be important [50]. We also restricted ourselves to the context of competition between two pathogens in a social host species. As the model was of a directly transmitted pathogen there is no transmission between individuals in separate groups. Infection via shared food sources or contact between individuals from different groups (e.g., during swarming [28]) would act to reduce population structure and therefore host density might become more important. However, further modelling would be required to demonstrate this while only empirical studies would be able to indicate the true relative importance of these different transmission routes in wild populations.

It is clear that many species are suffering strong population changes due to global change [17]. These changes might affect geographic range size [17], population size [18], population connectivity [51,52] or group size [53,54] to different degrees. The monitoring of these different aspects of population change, especially in bats, can often be difficult and may require further developments in monitoring to be effective, for example developing methods that use data from acoustic detectors [55–57]. Our results suggest that pathogen communities will respond differently depending on which traits are most strongly affected by global change. In short, species suffering reductions in groups size [53,54] are predicted to experience a decrease in pathogen richness in the long term and there is some evidence that this process is occurring [10,58]. Species that are experiencing an increase in group size [53] would be expected to gain new pathogen species and therefore pose a greater risk of being the source of a zoonotic disease. In contrast, species suffering a reduction in geographic range size [17] or a decrease in population size [18], without a corresponding decrease in group size, are expected to experience smaller changes in pathogen richness. Only by examining the mechanisms that control pathogen richness can we understand and predict these changes.

## Data accessibility

The implementation of the model is available as an R package on GitHub [36]. This can be found at https://github.com/timcdlucas/MetapopEpi. All other code and simulation output data is available on GitHub at https://github.com/timcdlucas/Abundance-Density-Manuscript.

## Author’s contributions

TCDL wrote the simulations and performed the analysis. TCDL, HMH and KEJ co-designed the study. TCDL drafted the manuscript. TCDL, HMH and KEJ all edited the manuscript and gave final approval for publication.

## Acknowledgements

We thank Andy Fenton and David Murrell for comments on the manuscript and help with the analytical model. This work, The Dynamic Drivers of Disease in Africa Consortium, NERC project no. NE-J001570-1 was funded with support from the Ecosystem Services for Poverty Alleviation Programme (ESPA). The ESPA programme is funded by the Department for International Development (DFID), the Economic and Social Research Council (ESRC) and the Natural Environment Research Council (NERC).

